# The Wolfram-like variant WFS1^E864K^ destabilizes MAM and compromises autophagy and mitophagy in human and mice

**DOI:** 10.1101/2023.11.16.567320

**Authors:** Simone Patergnani, Méghane S. Bataillard, Alberto Danese, Stacy Alves, Chantal Cazevieille, René Valéro, Lisbeth Tranebjærg, Tangui Maurice, Paolo Pinton, Benjamin Delprat, Elodie M. Richard

**Author notes:** These authors contributed equally (Simone Patergnani, Méghane S. Bataillard, Benjamin Delprat, Elodie M. Richard). To whom correspondence should be addressed: Elodie M. Richard Tel number: (33/0) 467 14 36 23 Fax number: Benjamin Delprat Tel number: (33/0) 467 14 36 23 Fax number: (33/0) 467 14 92 95.

## Abstract

Dominant variants in *WFS1*, a gene coding for the mitochondria-associated endoplasmic reticulum (ER) membrane (MAM) resident protein Wolframin, have been associated with Wolfram-like syndrome (WLS). *In vitro* and *in vivo*, WFS1 loss results in reduced ER to mitochondria calcium (Ca^2+^) transfer, mitochondrial dysfunction, and enhanced autophagy and mitophagy. However, in WLS pathological context, whether the mutant protein triggers the same cellular processes is unknown. Here, we show that, in human fibroblasts and murine neuronal cultures, WLS protein WFS1^E864K^ leads to decreases in mitochondria bioenergetics and Ca^2+^ uptake, deregulation of the mitochondrial quality system mechanisms, and alteration of the autophagic flux. Moreover, in the *Wfs1^E864K^* mouse, these alterations are concomitant with a decrease of MAM number. These findings reveal pathophysiological similarities between WS and WLS, highlighting the importance of WFS1 for MAM’s integrity and functionality. It may open new treatment perspectives, until now non-existent, for patients with WLS.

## Introduction

Wolframin, or WFS1, is a transmembrane protein, resident of the endoplasmic reticulum (ER) involved in cellular Ca^2+^ homeostasis and ER stress regulation ^1^. Variants in *WFS1*, the gene encoding the Wolframin, have been associated with Wolfram syndrome (OMIM #222300), a severe and rare neurodegenerative disease characterized by juvenile-onset diabetes, optic atrophy and deafness, as well as a wide diversity of endocrine and neurological alterations ^2,3^. Contrary to the recessive inheritance of WS, Wolfram-like syndrome (WLS, OMIM #614296) results from dominant *WFS1* variants and presents with the triad of symptoms that is progressive hearing loss, diabetes mellitus, and optic atrophy. Patients with WLS develop an overall milder phenotype, as there is few to no systemic manifestation leading to a shortened lifespan. However, vision and hearing capacities are severely impaired, with an earlier onset of the sensory deficits compared to WS patients. Even though no clear phenotype-genotype correlations can be made, the differences in symptomology between WS and WLS might rely on the type of variant and their inheritance ^4,5^. One can speculate that the dominant negative effect of missense variant associated with WLS might have increased toxicity in the sensory organs compared to the haploinsufficiency of biallelic nonsense variant associated with WS. Consistent with this hypothesis, in a mouse model, homozygous for the WLS *Wfs1^E864K^* allele ^6,7^, we recently showed the devastating effect of this missense variant on the inner ear ^8^. The homozygous mutant mice showed a profound post-natal hearing loss, at all tested frequencies, a collapse of the endocochlear potential and an alteration of the stria vascularis and neurosensory epithelium within their first month of age. While these effects might stem from an unbalanced ionic transfer due to a non-functional binding of WFS1 to the Na^+^ K^+^-ATPase ß1 subunit, we cannot exclude that other mechanisms are taking place in a tissue-specific manner, as suggested by a recent literature review ^5^.

Indeed, *WFS1* variant associated pathologies were first considered as resulting from an alteration of the ER stress response. As converging evidence, from independent studies, put light on the crucial role of WFS1 in Ca^2+^ dynamics at mitochondria-associated ER membranes (MAMs), leading eventually to mitochondrial bioenergetic deficits ^9–11^, they have recently been reclassified as “MAMopathies” ^12^. Numerous studies, using a variety of WFS1 knock-down or deficient models ranging from eukaryotic cells —HEK cells ^13^, rat primary cortical neurons ^9^, *Wfs1* null mouse derived neuronal cultures ^11^, human patients’ fibroblasts ^10^, patients’ hiPSC-derived neurons ^14^— to whole organisms —*wfs1* zebrafish ^15,16^— indicate that WFS1 is a key player for proper mitochondrial functionality, in link with a dysregulation of Ca^2+^ homeostasis via altered transfer between ER and mitochondria at the MAMs. In addition, in rat primary neuronal cultures ^9^, Wfs1 down-regulation impaired mitochondrial dynamics and led to mitophagy, a selective process that removed aged and damaged mitochondria. Similarly, in fibroblasts derived from WS patients, we have previously shown that loss of WFS1 increased not only the mitophagic processes but also the autophagic ones, both critical for proper mitochondrial functioning ^11^. Taken together, the dysregulations of Ca^2+^ homeostasis and mitochondrial functionality, due to an altered ER-mitochondria crosstalk, activate autophagic processes most likely responsible for the observed neurodegeneration in WS patients.

In contrast with WS pathophysiology, which has attracted attention in the recent years, little is known about the underlying molecular mechanisms of WLS and the impact of the WLS specific *WFS1* variants. In the present study, we put in light the molecular role of WFS1 in a pathophysiological context of WLS in the central nervous system of a mouse model of the disease ^8^, *Wfs1^E864K^*, and in patients’ fibroblasts ^7^, all harboring the missense variant _WFS1E864K._

## Results

In mouse and *in vitro* human models of WS, WFS1 deficiency has been associated with Ca^2+^ homeostasis alteration, mitochondrial dynamics impairment and decreased ER-mitochondria contacts, leading to increased autophagy and mitophagy ^10,11^. Patients with WLS present with less severe central deficits and one can question the impact of the disease-causing variant on the molecular pathways underlying the pathology.

To determine the functional impact of Wfs1^E864K^, a WLS-associated protein ^6,7^, we first investigated the mitochondrial respiration, using a Seahorse XFe analyzer, (Figure 1A) of neuronal cultures from the hippocampus and cortex of WT and homozygous mutant mice, *Wfs1^E864K^*, at post-natal day 33, age for which the neurosensorial deficits are at their maximal severity ^8^. We recorded a significant decrease of the basal (*p* = 0.005 and *p* < 0.0001; Figure 1B), ATP-related (*p* = 0.0018 and *p* = 0.0002; Figure 1C) and maximal (*p* = 0.0025 and *p* = 0.0004; Figure 1D) oxygen consumption rates of the neuronal cultures from the mutant mice, derived from both structures, compared to those of the cultures derived from WT mice. Likewise, we assessed directly on patients’ fibroblasts, heterozygous for the *WFS1^E864K^* variant, the impact of the WFS1^E864K^ mutant protein on the mitochondrial respiration (Figure 1E). Comparing control and patients’ fibroblasts, we measured a significant decrease of the basal (*p* = 0.019; Figure 1F), ATP-related (*p* = 0.0448; Figure 1G) and maximal (*p* = 0.0496; Figure 1H) oxygen consumption rates, as observed in mouse derived neuronal cultures.

**Figure 1.**
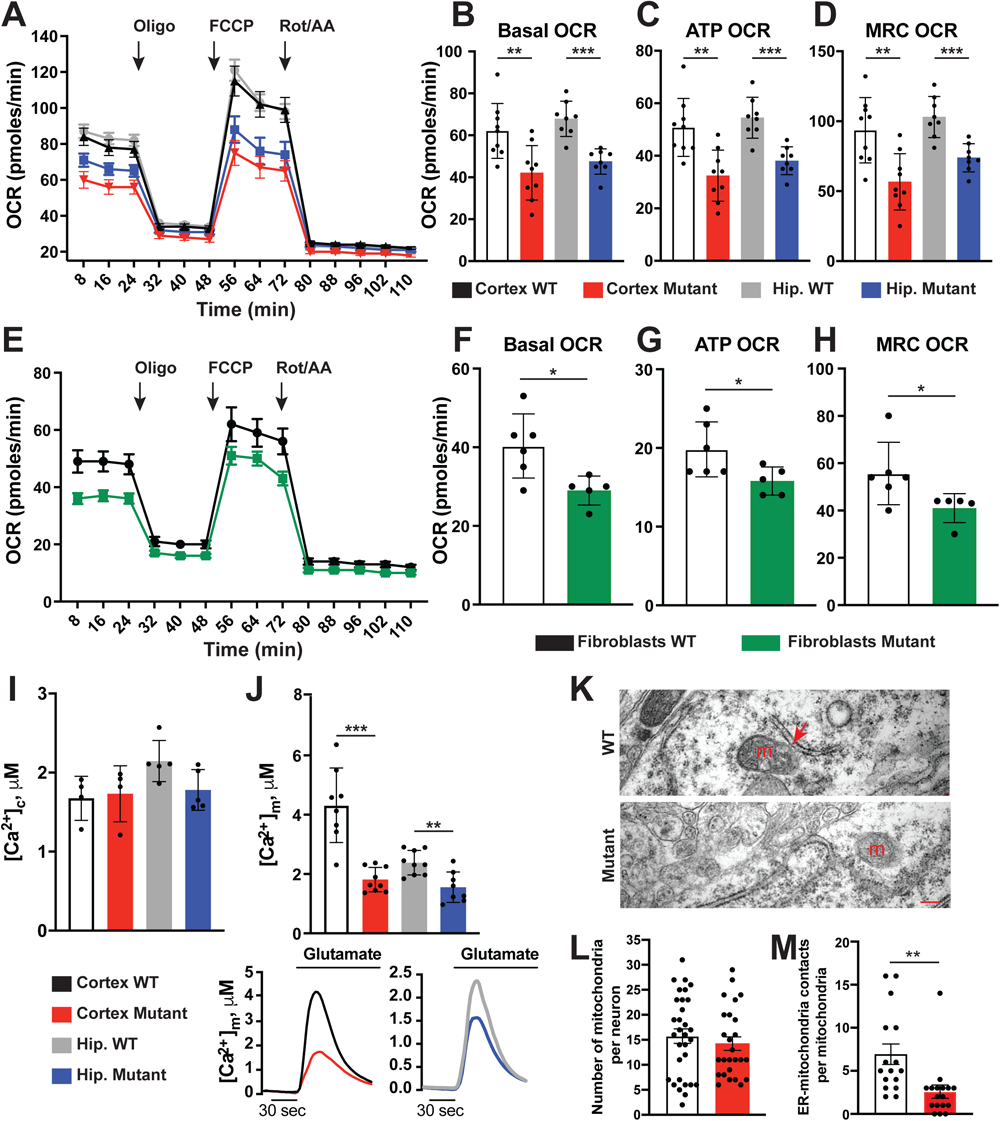
Wfs1^E864K^ protein leads to decreased mitochondrial respiration in patients’ fibroblasts and *Wfs1^E864K^* mutant mice, as well as altered Ca^2+^ transfer and a decrease of ER-mitochondria contacts in mice. (A-D) Oxygen consumption rate (OCR) traces of wild-type and mutant cortex and hippocampus (Hip.) neuronal cultures, expressed as picomoles of O_2_ per minute, under basal conditions and after the injection of oligomycin (1.5 μM), FCCP (1μM) and ROT/AA (1μM). Quantification of basal (B), ATP-related (C) and maximal (D) respiration rates, calculated from OCR traces and expressed as means ± SD. (E-H) Oxygen consumption rate (OCR) traces of control (WT) and patient fibroblasts (Mutant), expressed as picomoles of O_2_ per minute, under basal conditions and after the injection of oligomycin (1.5 μM), FCCP (1μM) and ROT/AA (1 μM). Quantifications of basal (F), ATP-related (G) and maximal (H) respiration rates, calculated from OCR traces and expressed as means ± SD. (I-J) Representative traces of aequorin-based measurements of neurons obtained from cortex and hippocampi of *Wfs1^WT^* (WT, cortex: black line, hippocampus: grey line) and *Wfs1^E864K^* (Mutant, cortex: black line, hippocampus: blue line) mice. Cytosolic Ca^2+^ (I) and mitochondrial Ca^2+^ (J) uptake measured after 10 μM glutamate stimulation. Quantification of cytosolic Ca^2+^ (I) and mitochondrial Ca^2+^ uptake (J), expressed as means ± SD, for hippocampus and cortex neuronal cultures of *Wfs1^WT^* (WT) and *Wfs1^E864K^* (mutant) mice. (K-M) Transmission electron micrographs of cortical neurons from three mutant and three WT mice. Contact between ER and outer membrane of a mitochondrion with a distance smaller than 30 nm is pointed by the red arrow. m: mitochondria, scale bar = 200 nm. Quantification of the number of mitochondria per neuron (L) and number of mitochondria in contact with ER (distance < 30 nm) (M). Data are expressed as means ± SEM. All tested animals were at 33 days-old at the time of experimentation. Per group, n = 8-9 for (B-D), n = 5-6 for (F-H), n = 4 for (I), n = 8-9 for (J) n = 24 (8 technical replicates and 3 biological replicates) for (L) and n = 16 (8 technical replicates and 2 biological replicates) for (M). Unpaired t-tests: * *p* < 0.05, ** *p* < 0.01, *** *p* < 0.001. Oligo, oligomycin; FCCP, carbonyl cyanide-4(trifluoromethoxy)phenylhydrazone; Rot/AA, rotenone/antimycin A.

In previous studies of WS models ^9–11^, the alteration of the mitochondrial functionality was associated with an impaired Ca^2+^ transfer between the ER and the mitochondria. Therefore, to assess, the Ca^2+^ dynamics, we performed aequorin experiments in neuronal cells stimulated with 10 μM glutamate. In response to glutamate, Ca^2+^ uptake from the mitochondria was significantly decreased (*p* < 0.0001 and *p* = 0.0022; Figure 1I). However, unlike the experiments on WS model ^10,11^, Ca^2+^ release from the ER into the cytosol was not affected in mutant neuronal cells compared to the WT cells (Figure 1J), pointing toward a mitochondrial defect rather than an ER alteration. Moreover, in the mutant and WT mice, we used transmission electron microscopy that allows a direct visualization of the MAMs, the contact zone between the ER and the mitochondria, to evaluate a potential structural alteration (Figure 1K). In the cortex of *Wfs1^E864K^* mice, the density of mitochondria per neuron was similar to that of the controls (Figure 1L). However, the number of contacts between the mitochondria and the ER was significantly reduced in the mutants (62% decrease in mutant cells compared to WT, *p* = 0.0042; Figure 1M), recapitulating what we observed in WS models ^11^. This structural defect, associated with the decrease of the Ca^2+^ uptake by the mitochondria, are more likely responsible for the dysfunction of the mitochondrial respiration.

Dysfunctional and damaged mitochondria are removed by mitophagy, a specialized cellular process that selectively remove these organelles by autophagy. Therefore, we investigated mitophagy, first in the cultured neuronal cells using two fluorescent probes to label the autolysosomes (lysotracker) and mitochondria (mitotracker) (Figure 2A). Confocal microscopy analysis revealed that in mutant neuronal cultured cells, the number of colocalizing dots was significantly increased compared to the number in the WT cells (*p* < 0.0001) (Figure 2B). Mitophagy is tightly linked to mitochondria biogenesis and turnover which allows the replacement of altered mitochondria and ultimately the maintenance of the pool of healthy mitochondria. Cells were transfected with MitoTimer ^17^, a fluorescent protein used to assess mitochondria turnover by labeling in green newly synthesized mitochondria while the fluorescence shifts to red in aged mitochondria (Figure 2C). Measures of the green and red fluorescent intensities in all cultured neuronal cells indicate that mitochondria in the mutant cells, from both hippocampal and cortical neurons, are older, and more likely less functional, that in the WT cells (*p* < 0.0001, *p* < 0.0001; Figure 2D).

**Figure 2.**
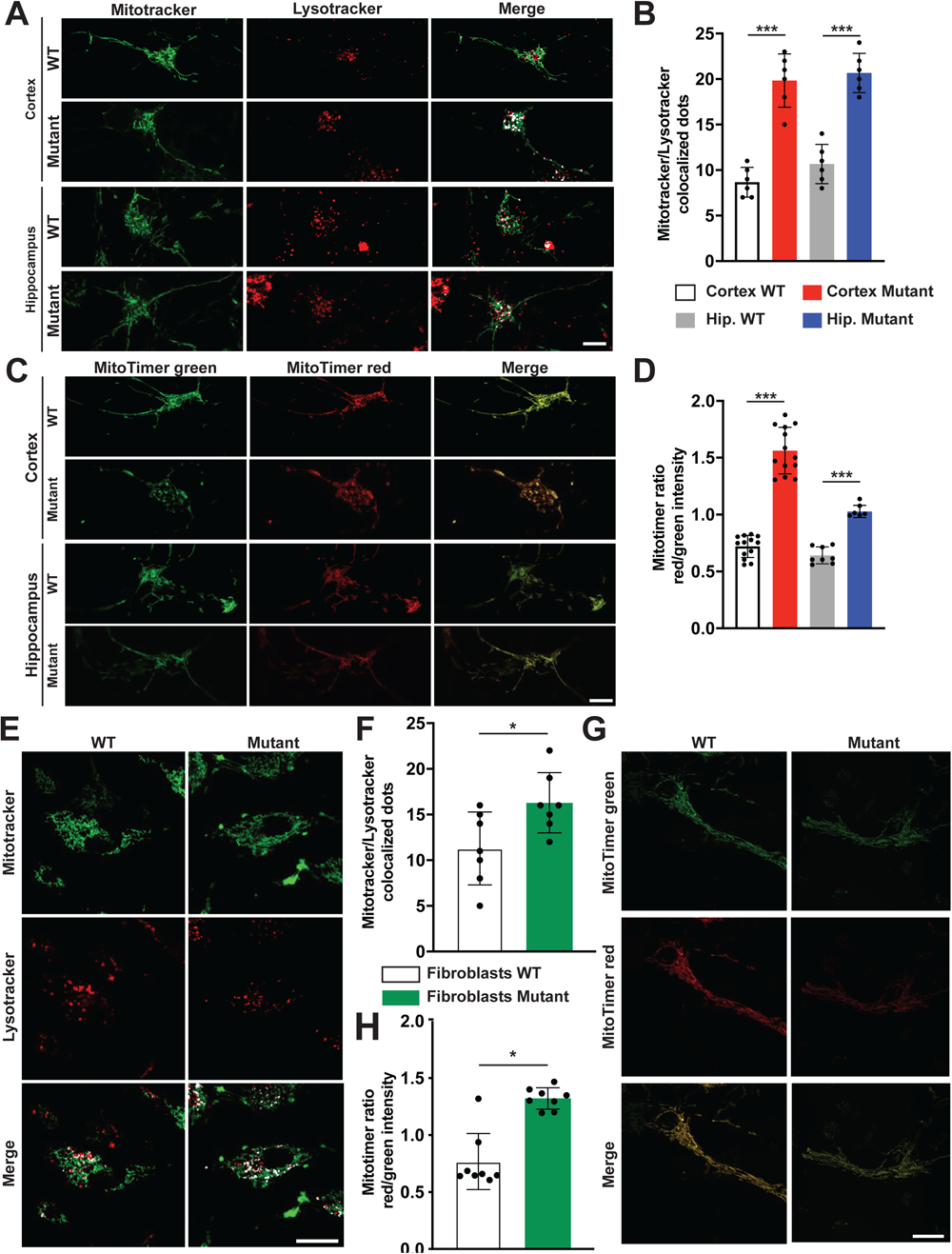
WFS1^E864K^ protein alters mitophagy and it is associated with an older mitochondrial population in fibroblasts from patient with WLS and neuronal cultures derived from WLS mouse model cortex and hippocampi. (A-B) Mitophagy activation assessment: (A) Confocal microscopy images of wild-type (WT) and mutant cortex and hippocampus (Hip) neuronal cultures labeled with mitochondrial (MitoTracker, green) and lysosomal (LysoTracker, red) markers and quantification of the colocalization dots (B). (C-D) MitoTimer measurements of mitochondrial age WT and mutant cortex and hippocampus neuronal cultures: (C) Representative confocal fluorescent images of the green, the red channels, and the merge. (D) Quantification of the red/green intensities ratio. (E-F) Mitophagy activation assessment: (E) Confocal microscopy images of fibroblasts labeled with mitochondrial (MitoTracker, green) and lysosomal (LysoTracker, red) markers and quantification of the colocalization dots (F). (G-H) Mitochondria turnover assessment using MitoTimer measurments in fibroblasts: (G) Representative epifluorescent images of the green, the red channels, and the merge. (H) Quantification of the red/green intensities ratio. Each value is the mean of n = 6 per condition in (B), n = 8-13 in (D), n = 7 in (F) and n = 8 in (H). Scale bar = 10 μm for all panels in (A) and (C), 20 μm for all panels in (E) and (G). Unpaired t-tests: * *p* < 0.05, *** *p* < 0.001.

In patients’ fibroblasts, even though less marked than in murine neuronal cells, we found an increase of the mitotracker-lysotracker colocalizing dots (*p* = 0.0252) in the mutant compared to the control human cells (Figure 2E-F). Similar to the study in mouse derived neuronal cells, MitoTimer experiments performed in patients’ fibroblasts lead to the conclusion of an older mitochondrial population in cells carrying the *WFS1^E864K^* variant compared to controls (*p* < 0.0001; Figure 2G-H).

In WS, the absence of WFS1 is associated with an enhanced autophagy ^11^ linked to an unbalanced Ca^2+^ homeostasis. This dysregulation could be observed in the primary culture of hippocampal and cortical neurons. Cultured neurons were collected for immunoblot using the specific autophagic markers microtubule-associated proteins 1A/1B light chain 3 (LC3) and P62, respectively used for the detection of the early and late steps of the autophagy process (Figure 3A). Interestingly, we found an increased level of LC3-II (Figure 3B), the cleaved form of LC3 related to the activation of the autophagic process, accompanied by a concomitant increase of P62 level (Figure 3C) in the mutant cultured neurons (*p* = 0.0004, *p* < 0.0001 for LC3-II; *p* < 0.0001, *p* < 0.0001 for P62). Similar increases were observed in fibroblasts from WLS patients by western blot analysis (*p* = 0.0074 for LC3-II, *p* = 0.0039 for P62; Figure 3D-F). A simultaneous increase of both the cleaved form of LC3 and of P62 has been frequently associated to a dysregulation of the autophagic flux, in which the autophagosomes are continuously created (LC3-II increase) but they struggle to be degraded (P62 accumulation) ^18^. We investigated further these autophagy dynamics using fibroblast cells transfected with a green fluorescent protein (GFP)-LC3 construct (GFP-LC3^vac^, ^19^), which allows the visualization and quantification of the autophagosomes as punctate structures (Figure 3G). The elevated amount of LC3-II measured by western blot in mutant cells was confirmed by fluorescent microscopy analysis of GFP-LC3 transfected fibroblasts (Figure 3G-H), where green autophagosomal puncta were more abundant than in WT cells (*p* < 0.0001). To discriminate whether the impact of the mutant WFS1^E864K^ protein on the autophagic processes is due to an activation of the autophagic flux or an inhibition of the autophagosome-lysosome fusion, we performed the same microscopy analysis with cells treated with bafilomycin A1 (BafA1), an antibiotic which impairs the autophagosome-lysosome fusion and autolysosome acidification, the late steps in the autophagic process, blocking the autophagic flux. Fluorescent microscopy analysis of the punctate structures after BafA1 treatment suggests that mutant WFS1 protein, rather than activating a sustained autophagosome formation process, alters and slows down the autophagy flux. Indeed, post-treatment, the number of green puncta per cell increases in both conditions, WT and mutant fibroblasts, however the increase in the mutant is comparatively lower than in the WT (+ 30% in the mutant cells *vs*. + 300% in the WT cells) (Figure 3G-H). Overall, these data clearly indicated that the mutant protein might interfere with the autophagy flux, compromising the autolysosomal processes.

**Figure 3.**
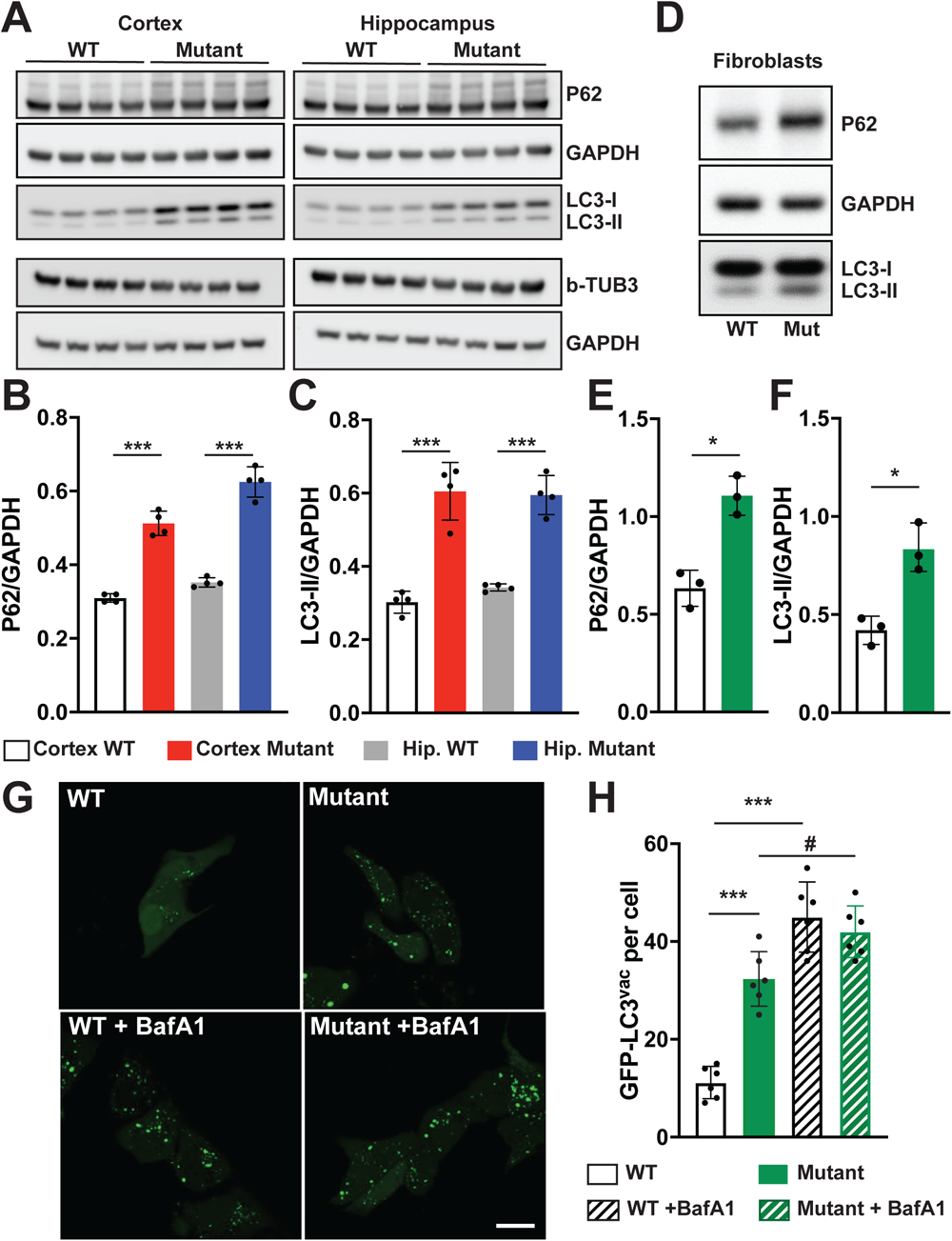
WFS1^E864K^ protein induces an altered autophagy flux in mouse neurons and fibroblasts. (A-C) Autophagy in neuronal cultured derived from *Wfs1^E864K^* mice: (A) Representative immunoblot showing the increase of P62 and the autophagic form of LC3 (LC3-II) in neurons cultured from the cortex and hippocampus (Hip) of *Wfs1^WT^* (WT) and *Wfs1^E864K^* (Mutant) mice. The purity of neuronal cultures was assessed with immunoblot against the neuronal marker β3-Tubulin (b-TUB3). GAPDH was used as loading control. Densitometry ratios of P62 and LC3-II over GAPDH are shown in (B) and (C). (D-F) Autophagy in patient’s fibroblasts: (D) Representative immunoblot showing the increase of P62 and LC3 lipidation in fibroblasts from control (WT) and patient harboring the WFS1^E864K^ variant (Mut). GAPDH was used as loading control. Densitometry ratios of P62 and LC3-II over GAPDH are shown in (E) and (F). (G) Representative images of fluorescence microscopy acquisition of fibroblasts transfected with GFP-LC3^vac^ plasmid. To mimic a blockage of the autophagic flux, 100 nM bafilomycin A1 (BafA1) was added for 2 h. (H) Quantification of LC3-autophagic dot per cell. Each value is the mean of n = 4 per condition in (B, C), n = 3 in (E, F), and n = 6 in (H). Scale bar = 20 μm. Unpaired t-tests for (B-F): * *p* < 0.05, *** P < 0.001. Two-way ANOVA followed by Tukey’s multiple comparisons test for (H): P < 0.0001 for the genotype, P < 0.0001 for the BafA1 treatment, *** P < 0.001 vs. WT, # P < 0.05 vs. Mutant.

## Discussion

WFS1 variants associated disorders encompass a spectrum of rare pathologies ^20^, recently termed “wolframinopathies” by de Muijnck and coll. ^5^. WS is the most common one even though its worldwide prevalence is estimated to approximately 1 in 770,000 ^2^. *WFS1* biallelic recessive variants are responsible for Wolfram syndrome type 1. Given the resemblance of WS symptoms with mitochondrial diseases, mitochondrial functionality was studied in different models of the syndrome and highlighted the key role of WFS1 in mitochondrial bioenergetics ^9,11,13–16^. However, *WFS1* expression was either abolished or knocked down in these models, leading to a low level to the total absence of WFS1 protein. In the context of WLS, the mutant protein is expressed, along the WT protein, so the molecular mechanism(s) underlying the pathology might be different.

In our study, we determined that WFS1^E864K^ mutant protein is associated with mitochondrial bioenergetic deficits, both in murine neuronal cells and fibroblasts derived from WLS patients. These results were comparable to the one obtained in similar cellular models of WFS1 loss ^10,11^. One explanation for altered energy production is a dysregulated calcium homeostasis, as observed in various WS models. Contrary to what we previously reported in WS cellular models, cytosolic Ca^2+^ was not different between WT and mutant cells suggesting a normal ER efflux of Ca^2+^. However, mitochondrial Ca^2+^ uptake was decreased in mutant neuronal cell line, pointing toward an alteration at the mitochondria level rather than from the ER. This reduced uptake, associated with normal intracellular Ca^2+^ dynamics, can be explained by a decreased interaction between these 2 organelles. Indeed, despite a similar number of mitochondria per neuron, the number of contacts between the mitochondria and the ER, defining the number of MAMs, is lower in mutant cortical neurons compared to WT cells. At the ER-mitochondria cleft, Ca^2+^ stored in the ER is released by the inositol 1,4,5-trisphosphate receptors (IP3R), then transported to the voltage-dependent anion channel (VDAC1) by the glucose regulatory protein 75 (GRP75), the mitochondrial calcium uniporter (MCU) eventually channeling the cations to the mitochondrial matrix ^21^. We specifically looked at contacts with a distance of less than or equal to 30 nm, the width necessary to allow correct assembly of the IP_3_R-GRP75-VDAC1 complex, responsible for Ca^2+^ flux ^22^. Consequently, we can speculate that this diminished interaction can adversely affect Ca^2+^ transfer and homeostasis, even though we cannot exclude a change in mitochondria membrane potential or in expression level of mitochondria Ca^2+^ uptake machinery ^23^.

Samples obtained from mutated mice as well as fibroblasts from WLS patients displayed an aged population of mitochondria, which can correlate with the bioenergetic deficits measured. Having found an affected mitophagy machinery in these cells, this data suggest that aged and not functional mitochondria are not removed correctly. To date, mitophagy initiation and progression exact mechanisms remain unclear. Converging lines of evidence support a strong relationship between MAM and mitophagy ^24^. Thus, the altered MAM integrity and function in WLS context might be leading to the activation of mitophagy and consequently the alteration of mitochondrial functions. Likewise, the increase of autolysosomal processes in mutant cells might be explained by the deficient ER-mitochondria coupling in these cells. The origin of the autophagosome membrane is still under debate, however, ample evidence suggest that autophagy begins at the MAMs ^25^. They would play a dual role as a platform for autophagy related protein to perform their biological function and as a modulator of the Ca^2+^ homeostasis and lipid metabolism which are both key for autophagosome development ^24,25^. Indeed, unbalanced Ca^2+^ homeostasis between ER and mitochondria has been shown to activate pro-survival autophagy at MAM level ^26,27^. A decrease of Ca^2+^ transfer from the ER to the mitochondria was shown to activate AMPK translocation to MAMs which would activate autophagy processes via BECN protein ^28^. Thus, the increase of autophagy markers in *WFS1^E864K^* fibroblasts and neuronal cells may arise from the observed MAM alteration. We showed in WS patients’ fibroblasts that the absence of WFS1 activated both mitophagy and autophagy processes ^11^. However, contrary to what was observed in WFS1 null fibroblasts, when the autophagy flux is blocked using BafA1, the relative rise of autophagosomes is less in the mutant than in the WT fibroblasts, suggesting an alteration of the autophagic machinery of these cells in basal conditions. A compromised autophagy is detrimental for neuronal cells, often time associated with neurodegeneration ^29^, as reported not only in the pathophysiological context of WS ^9,15^ but also other neurodegenerative disorders ^30^. The attenuated dysregulation of autophagy might explain why no degeneration has been reported so far in WLS patients ^5^. However, the rarity of the disease and the underdiagnose of the population might have hampered an extended characterization of the symptomology of WLS, more particularly in aged patients.

Akin what we reported in WS models ^10,11^, overall, the decreased number of MAMs is triggering an alteration of the cellular bioenergetics and a deregulation of ER-mitochondria crosstalk. These cellular and molecular abnormalities have been reported for neurodegenerative disorders such as Alzheimer disease, Parkinson disease, Huntington disease, and amyotrophic lateral sclerosis, among others ^24,31–33^. In addition, new studies highlighted the potential role of WFS1 itself in central nervous system neurodegeneration and more particularly Alzheimer disease ^9,34^. Thus, even though the severe vestibular syndrome of *Wfs1^E864K^*mice is preventing us to explore their potential behavioral deficits, in light of these shared common mechanisms underlying WS, WLS and the other neurodegenerative disorders, it would be of great interest to look at aged mice to investigate potential neurodegenerative processes.

While the importance of WFS1 for proper MAM function has been well established using *in vitro* and *in vivo* models, its role(s) in the structure of the MAM remain unknown. In mouse and human, WFS1 regulates the number of contacts between the ER and the mitochondria ^10,11^. One explanation could be that WFS1 modulates tethering proteins, warrants of the structural integrity of the MAMs. Indeed, the combination of optic atrophy and hearing loss is the clinical hallmark of WLS, but also dominant optic atrophy and deafness (DOAD, OMIM 125250). Variants in *OPA1*, a gene encoding the mitochondrial dynamin-like GTPase (OPA1) protein, are mainly responsible for the disease ^35^. This protein, anchored to the mitochondrial inner membrane facing the inter-membrane space, plays a critical role in mitochondria, as a link between the mitochondrial structure and bioenergetic functions, regulating mitochondrial fusion ^36^. This fusion process of mitochondrial membranes is promoted by the binding of OPA1 and the mitochondria-bound Mitofusin 2 protein (MFN2), a key tethering protein of the MAMs (for review ^37^). In the absence of WFS1, in rat and patients’ fibroblasts, no change of *MFN2* expression level was reported ^9,10^. However, due to the similarities between DOAD and WSL symptoms, one can ask if the Ca^2+^ homeostasis and MAM alteration in WSL might be due to tethering abnormalities rather than stemming primarily from IP_3_R-GRP75-VDAC1 complex function alteration. In addition, WFS1 is interacting with several MAM resident proteins such as Sigma-1 receptor (S1R) ^11^, neuronal calcium sensor 1 (NCS1) ^10^ and VDAC1 itself ^14^, which all have a role in MAM structure as well as function. Even though we cannot qualify VDAC1 role as classical tethering, it works as a contact site spacing or filling function through its interaction with GRP75 and IP_3_R ^14^. S1R and NCS1 stabilize MAMs by interacting with IP_3_R and/or VDAC1 ^25,38^. Variants in WFS1 might impact the binding to its protein partners leading to aberrant MAM structure and function.

Overall, in murine neuronal cells and patients’ fibroblasts, our findings indicate that WFS1^E864K^ protein leads to aberrant Ca^2+^ homeostasis, mitochondrial bioenergetics deficits, deregulation of both mitophagy and autophagy, most likely stemming from MAM alterations. Our findings suggest considering the use of the same therapeutic targets as for WS in a WLS context, combined with autophagy inducers to ameliorate the outcome, thus opening up treatment perspectives that were previously non-existent.

## Materials and Methods

### Fibroblasts culture

This study followed the tenets of the Declaration of Helsinki. Primary cultures of human dermal fibroblasts were established from skin biopsies taken after obtaining informed consent from one individual without diabetes mellitus and one affected patient carrying the c.2590G>A (E864K) variant in *WFS1* gene, as previously described ^39^. Fibroblasts from these skin explants were cultured in Dulbecco modified Eagle’s medium (DMEM) supplemented with glucose (4.5 g l^−1^), 2mM l-glutamine, 10% (v/v) fetal bovine serum (FBS) (Gibco) and 1% penicillin– streptomycin. All the experiments were performed on fibroblasts between tissue culture passages 4 and 12.

Transfection with the autophagy reporter GFP-LC3 and the indicator of mitochondrial turnover pTRE-tight-MITO TIMER was performed using Lipofectamine LTX&PLUS (Life Technologies, 15338-100).

### Mouse breeding

The *Wfs1^E864K^* mice were generated as previously described ^8^. Animal care and experimentation were authorized by the National Ethic Committee (Paris) and carried out in strict adherence to the European Union Directive 2010/63 and ARRIVE guidelines ^40^. Mice were housed in cages with free access to water and food. The rooms were temperature-controlled, lit on a 12:12 h light/dark cycle, lights on at 7:00 h. All experiments were performed during the light cycle, at the same time.

### Isolation of neurons from Wfs1^WT^ and Wfs1^E864K^ hippocampi and cortex

Primary neuronal cultures were obtained from brains of adult mice as reported in ^11^. Briefly, after isolation from the brains, hippocampi and cortex were first mechanically and then enzymatically dissociated at 37 °C for 15 minutes in a solution of 0.25% (wt/vol) trypsin and 10 μg/ml DNase. After filtration with a 70 μM pores cell strainer, the cellular homogenates obtained were cultured in neuronal culture medium Neurobasal (Thermo Fisher Scientific, catalog number: 21103-049) enriched with B27 supplement (Thermo Fisher Scientific, catalog number: 17504-044), 1% penicillin–streptomycin and 0.5 mM GlutamaX. To avoid the proliferation of non-neuronal cells, cytosine arabinoside 1 µM was added to the cultures after 24 hours from the plating. Immunoblot assays against specific neuronal markers β3-Tubulin (Cell Signaling Technology, 5568) permitted to assess the purity of cell cultures.

### Aequorin measurements

Cells were seeded on 13 mm glass coverslips and transduced with adenoviral construct encoding a cytosolic and mitochondria-targeted aequorin. 36 hours post-infection, cells were reconstituted for 2 hours with 5 µM coelenterazine wt (Santa Cruz Biotechnology) in DMEM high glucose supplemented of 0.1% FBS and transferred to the perfusion chamber. Measurements were performed in Krebs-Ringer buffer (135 mM NaCl, 5 mM KCl, 1 mM MgSO_4_, 0.4 mM KH_2_PO_4_, 20 mM HEPES, pH 7.4) supplemented with 5 mM glucose and 1 mM CaCl_2_. Calcium (Ca^2+^) release from the intracellular stores and the consequent cytosolic and mitochondrial Ca^2+^ accumulation was induced by adding the Ca^2+^-mobilizating agent glutamate. After this stimulation, cells were lysed with 100 μM digitonin in a hypotonic Ca^2+^-rich solution (10 mM CaCl_2_ in H_2_O), to discharge the remaining aequorin pool. Finally, the light signal detected was collected and calibrated into Ca^2+^ values by using a specific algorithm, as described in ^41^.

### MAM analysis (Transmission Electron Microscopy)

Morphological changes were investigated using TEM analyses. *Wfs1^WT^*and *Wfs1^E864K^*homozygous mice were perfused intracardially with a 4% paraformaldehyde solution at room temperature (n=3 mice per genotype). Small fragments (about 1 mm^3^) of the cortex from the brain of each mouse were extracted and immersed in a solution of 2.5% glutaraldehyde in PHEM buffer (1X, pH 7.4) overnight at 4°C. They were then rinsed in PHEM buffer and post-fixed in a 0.5% osmic acid + 0.8% potassium Hexacyanoferrate trihydrate for 2 h at dark and room temperature. After two rinses in PHEM buffer, the cells were dehydrated in a graded series of ethanol solutions (30-100%). The tissues were embedded in EmBed 812 using an Automated Microwave Tissue Processor for Electronic Microscopy (Leica, Wetzlar, Germany). Thin sections (70 nm; Leica-Reichert Ultracut E) were collected at different levels of each block. These sections were counterstained with uranyl acetate 1.5% in 70% Ethanol and lead citrate and observed using a Tecnai F20 transmission electron microscope (FEI company, Hillsboro, Oregon, USA).

To evaluate the frequency of contact between the mitochondria and the ER in the different neurons, we photographed at 5,800 x magnification, all the cytoplasmic area from 8 different neurons in each of 6 samples. with a Tecnai F20 transmission electron microscope at 120 KV. From these images, we counted the number of mitochondria. We then randomly photographed at 25,500 x magnification, 8 different cytoplasmic area, containing mitochondria, from the previous neurons. In each image, we counted the number of mitochondria and calculated the proportion of mitochondria in close contact with ER (< 30 nm).

### Autophagy analyses

Primary antibodies used for the detection of the autophagic mechanisms by immunoblot were anti-LC3B antibody (Merck Group, L7543) and anti-SQSTM1/p62 (Merck Group, P0067). An anti-beta-actin (Merck Group, A1978) antibody was used as loading marker. Autophagy was also detected by fluorescent microscopy by transfecting cells with a GFP-LC3 construct. Images were taken on an Olympus FV3000 confocal microscope (Olympus Corporation, Tokyo, Japan) equipped of a 63x oil immersion objective (N.A. 1.4). The number of GFP-LC3 dots was automatically counted by using ImageJ (Fiji) software. Where indicated cells were treated with bafilomycin A1 (BafA1), a potent inhibitor of the lysosomal V-ATPase, to mimic the inhibition of autophagosome-lysosome fusion, as reported in ^42^.

### Mitophagy analyses

Cells were seeded on 24 mm glass coverslips and loaded with the mitochondrial and lysosomal fluorescent probes, MitoTracker Green FM (Life Technologies) and LysoTracker Red DND-99 (Life Technologies), respectively, for 15 minutes. Next cells were taken by using a FV3000 Olympus confocal microscope with a 63x oil immersion objective (N.A. 1.4). Images were background subtracted and the co-localization rates were measured by using the co-localization plugin “Colocalization Thresholds” available with Fiji software.

### Statistical analyses

Data are expressed as mean ± SD. Statistical significance between groups was determined by unpaired Student’s t test or 2way ANOVA followed by Tukey’s multiple comparisons test. The levels of statistical significance considered were: * P < 0.05, ** P < 0.01, and *** P < 0.001. Statistical analyses were performed using the Prism v.7.0 software (GraphPad, San Diego, CA).

#### Abbreviation

BafA1: Bafilomycin
ER: endoplasmic reticulum
GRP75: glucose regulatory protein 75
IP_3_R: inositol 1,4,5-trisphosphate receptor
MAM: mitochondria-associated endoplasmic reticulum membrane
MCU: mitochondrial calcium uniporter
MFN2: Mitofusin 2 protein
VDAC1: voltage-dependent anion channel
WLS: Wolfram-like syndrome
WS: Wolfram syndrome
WT: wild-type

## Acknowledgments

We thank the patient and their family for their contribution to the study, Marc Thiry (University of Liege, Belgium) for his unvaluable advice for TEM analysis. We thank the breeding facility from the University of Montpellier (RAM-CECEMA) for their help in animal breeding and handling. We thank the Electronic Microscopy facilities of the Institut des Neurosciences de Montpellier for TEM acquisition. P.P. is grateful to C. degli Scrovegni for her continuous support.

## References

1. Fonseca SG, Fukuma M, Lipson KL, Nguyen LX, Allen JR, Oka Y, Urano F. WFS1 is a novel component of the unfolded protein response and maintains homeostasis of the endoplasmic reticulum in pancreatic beta-cells. J Biol Chem 2005; 280:39609–15.

2. Barrett TG, Bundey SE, Macleod AF. Neurodegeneration and diabetes: UK nationwide study of Wolfram (DIDMOAD) syndrome. Lancet 1995; 346:1458–63.

3. Rigoli L, Bramanti P, Di Bella C, De Luca F. Genetic and clinical aspects of Wolfram syndrome 1, a severe neurodegenerative disease. Pediatr Res 2018; 83:921–9.

4. de Heredia ML, Clèries R, Nunes V. Genotypic classification of patients with Wolfram syndrome: insights into the natural history of the disease and correlation with phenotype. Genet Med 2013; 15:497–506.

5. de Muijnck C, Brink JBT, Bergen AA, Boon CJF, van Genderen MM. Delineating Wolfram-like syndrome: A systematic review and discussion of the WFS1-associated disease spectrum. Surv Ophthalmol 2023; 68:641–54.

6. Eiberg H, Hansen L, Kjer B, Hansen T, Pedersen O, Bille M, Rosenberg T, Tranebjaerg L. Autosomal dominant optic atrophy associated with hearing impairment and impaired glucose regulation caused by a missense mutation in the WFS1 gene. J Med Genet 2006; 43:435–40.

7. Valéro R, Bannwarth S, Roman S, Paquis-Flucklinger V, Vialettes B. Autosomal dominant transmission of diabetes and congenital hearing impairment secondary to a missense mutation in the WFS1 gene. Diabet Med 2008; 25:657–61.

8. Richard EM, Brun E, Korchagina J, Crouzier L, Affortit C, Alves S, Cazevieille C, Mausset-Bonnefont A-L, Lenoir M, Puel J-L, et al. Wfs1E864K knock-in mice illuminate the fundamental role of Wfs1 in endocochlear potential production. Cell Death Dis 2023; 14:387.

9. Cagalinec M, Liiv M, Hodurova Z, Hickey MA, Vaarmann A, Mandel M, Zeb A, Choubey V, Kuum M, Safiulina D, et al. Role of Mitochondrial Dynamics in Neuronal Development: Mechanism for Wolfram Syndrome. PLoS Biol 2016; 14:e1002511.

10. Angebault C, Fauconnier J, Patergnani S, Rieusset J, Danese A, Affortit CA, Jagodzinska J, Mégy C, Quiles M, Cazevieille C, et al. ER-mitochondria cross-talk is regulated by the Ca2+ sensor NCS1 and is impaired in Wolfram syndrome. Sci Signal 2018; 11.

11. Crouzier L, Danese A, Yasui Y, Richard EM, Liévens J-C, Patergnani S, Couly S, Diez C, Denus M, Cubedo N, et al. Activation of the sigma-1 receptor chaperone alleviates symptoms of Wolfram syndrome in preclinical models. Sci Transl Med 2022; 14:eabh3763.

12. Delprat B, Maurice T, Delettre C. Wolfram syndrome: MAMs’ connection? Cell Death Dis 2018; 9:364.

13. Kõks S, Overall RW, Ivask M, Soomets U, Guha M, Vasar E, Fernandes C, Schalkwyk LC. Silencing of the WFS1 gene in HEK cells induces pathways related to neurodegeneration and mitochondrial damage. Physiol Genomics 2013; 45:182–90.

14. Zatyka M, Rosenstock TR, Sun C, Palhegyi AM, Hughes GW, Lara-Reyna S, Astuti D, di Maio A, Sciauvaud A, Korsgen ME, et al. Depletion of WFS1 compromises mitochondrial function in hiPSC-derived neuronal models of Wolfram syndrome. Stem Cell Reports 2023; 18:1090–106.

15. Crouzier L, Richard EM, Diez C, Alzaeem H, Denus M, Cubedo N, Delaunay T, Glendenning E, Baxendale S, Liévens J-C, et al. Morphological, behavioral and cellular analyses revealed different phenotypes in Wolfram syndrome wfs1a and wfs1b zebrafish mutant lines. Hum Mol Genet 2022;: ddac065.

16. Crouzier L, Richard EM, Diez C, Denus M, Peyrel A, Alzaeem H, Cubedo N, Delaunay T, Maurice T, Delprat B. NCS1 overexpression restored mitochondrial activity and behavioral alterations in a zebrafish model of Wolfram syndrome. Mol Ther Methods Clin Dev 2022; 27:295–308.

17. Hernandez G, Thornton C, Stotland A, Lui D, Sin J, Ramil J, Magee N, Andres A, Quarato G, Carreira RS, et al. MitoTimer. Autophagy 2013; 9:1852–61.

18. Klionsky DJ, Abdel-Aziz AK, Abdelfatah S, Abdellatif M, Abdoli A, Abel S, Abeliovich H, Abildgaard MH, Abudu YP, Acevedo-Arozena A, et al. Guidelines for the use and interpretation of assays for monitoring autophagy (4th edition)1. Autophagy 2021; 17:1– 382.

19. Patergnani S, Marchi S, Rimessi A, Bonora M, Giorgi C, Mehta KD, Pinton P. PRKCB/protein kinase C, beta and the mitochondrial axis as key regulators of autophagy. Autophagy 2013; 9:1367–85.

20. Tranebjærg L, Barrett T, Rendtorff ND. WFS1 Wolfram Syndrome Spectrum Disorder [Internet]. In: Adam MP, Everman DB, Mirzaa GM, Pagon RA, Wallace SE, Bean LJ, Gripp KW, Amemiya A, editors. GeneReviews®. Seattle (WA): University of Washington, Seattle; 1993 [cited 2022 Sep 1]. Available from: http://www.ncbi.nlm.nih.gov/books/NBK4144/

21. Giorgi C, Marchi S, Pinton P. The machineries, regulation and cellular functions of mitochondrial calcium. Nat Rev Mol Cell Biol 2018; 19:713–30.

22. Giacomello M, Pellegrini L. The coming of age of the mitochondria-ER contact: a matter of thickness. Cell Death Differ 2016; 23:1417–27.

23. Rossi A, Pizzo P, Filadi R. Calcium, mitochondria and cell metabolism: A functional triangle in bioenergetics. Biochim Biophys Acta Mol Cell Res 2019; 1866:1068–78.

24. Liu J, Yang J. Mitochondria-associated membranes: A hub for neurodegenerative diseases. Biomedicine & Pharmacotherapy 2022; 149:112890.

25. Yang M, Li C, Yang S, Xiao Y, Xiong X, Chen W, Zhao H, Zhang Q, Han Y, Sun L. Mitochondria-Associated ER Membranes – The Origin Site of Autophagy. Front Cell Dev Biol 2020; 8:595.

26. Cárdenas C, Miller RA, Smith I, Bui T, Molgó J, Müller M, Vais H, Cheung K-H, Yang J, Parker I, et al. Essential regulation of cell bioenergetics by constitutive InsP3 receptor Ca2+ transfer to mitochondria. Cell 2010; 142:270–83.

27. Missiroli S, Bonora M, Patergnani S, Poletti F, Perrone M, Gafà R, Magri E, Raimondi A, Lanza G, Tacchetti C, et al. PML at Mitochondria-Associated Membranes Is Critical for the Repression of Autophagy and Cancer Development. Cell Rep 2016; 16:2415– 27.

28. Ahumada-Castro U, Silva-Pavez E, Lovy A, Pardo E, Molgό J, Cárdenas C. MTOR-independent autophagy induced by interrupted endoplasmic reticulum-mitochondrial Ca2+ communication: a dead end in cancer cells. Autophagy 2018; 15:358–61.

29. Aman Y, Schmauck-Medina T, Hansen M, Morimoto RI, Simon AK, Bjedov I, Palikaras K, Simonsen A, Johansen T, Tavernarakis N, et al. Autophagy in healthy aging and disease. Nat Aging 2021; 1:634–50.

30. Giorgi C, Bouhamida E, Danese A, Previati M, Pinton P, Patergnani S. Relevance of Autophagy and Mitophagy Dynamics and Markers in Neurodegenerative Diseases. Biomedicines 2021; 9:149.

31. Fernandes T, Resende R, Silva DF, Marques AP, Santos AE, Cardoso SM, Domingues MR, Moreira PI, Pereira CF. Structural and Functional Alterations in Mitochondria-Associated Membranes (MAMs) and in Mitochondria Activate Stress Response Mechanisms in an In Vitro Model of Alzheimer’s Disease. Biomedicines 2021; 9:881.

32. Johri A, Chandra A. Connection Lost, MAM: Errors in ER–Mitochondria Connections in Neurodegenerative Diseases. Brain Sci 2021; 11:1437.

33. Degechisa ST, Dabi YT, Gizaw ST. The mitochondrial associated endoplasmic reticulum membranes: A platform for the pathogenesis of inflammation-mediated metabolic diseases. Immunity, Inflammation and Disease 2022; 10:e647.

34. Li L, Venkataraman L, Chen S, Fu H. Function of WFS1 and WFS2 in the Central Nervous System: Implications for Wolfram Syndrome and Alzheimer’s Disease. Neurosci Biobehav Rev 2020; 118:775–83.

35. Delettre C, Lenaers G, Griffoin J-M, Gigarel N, Lorenzo C, Belenguer P, Pelloquin L, Grosgeorge J, Turc-Carel C, Perret E, et al. Nuclear gene OPA1, encoding a mitochondrial dynamin-related protein, is mutated in dominant optic atrophy. Nat Genet 2000; 26:207–10.

36. Del Dotto V, Carelli V. Dominant Optic Atrophy (DOA): Modeling the Kaleidoscopic Roles of OPA1 in Mitochondrial Homeostasis. Front Neurol 2021; 12:681326.

37. Zaman M, Shutt TE. The Role of Impaired Mitochondrial Dynamics in MFN2-Mediated Pathology. Frontiers in Cell and Developmental Biology [Internet] 2022 [cited 2023 Sep 1]; 10. Available from: 10.3389/fcell.2022.858286

38. Su T-P, Hayashi T, Maurice T, Buch S, Ruoho AE. The sigma-1 receptor chaperone as an inter-organelle signaling modulator. Trends Pharmacol Sci 2010; 31:557–66.

39. Angebault C, Gueguen N, Desquiret-Dumas V, Chevrollier A, Guillet V, Verny C, Cassereau J, Ferre M, Milea D, Amati-Bonneau P, et al. Idebenone increases mitochondrial complex I activity in fibroblasts from LHON patients while producing contradictory effects on respiration. BMC Res Notes 2011; 4:557.

40. Kilkenny C, Browne WJ, Cuthill IC, Emerson M, Altman DG. Improving bioscience research reporting: The ARRIVE guidelines for reporting animal research. J Pharmacol Pharmacother 2010; 1:94–9.

41. Bonora M, Giorgi C, Bononi A, Marchi S, Patergnani S, Rimessi A, Rizzuto R, Pinton P. Subcellular calcium measurements in mammalian cells using jellyfish photoprotein aequorin-based probes. Nat Protoc 2013; 8:2105–18.

42. Patergnani S, Bonora M, Ingusci S, Previati M, Marchi S, Zucchini S, Perrone M, Wieckowski MR, Castellazzi M, Pugliatti M, et al. Antipsychotic drugs counteract autophagy and mitophagy in multiple sclerosis. Proc Natl Acad Sci U S A 2021; 118:e2020078118.

